# A first pangenomic framework for globe artichoke supports SNP-based varietal fingerprinting

**DOI:** 10.64898/2026.06.25.734495

**Authors:** Ezio Portis, Edoardo Vergnano, Luciana Gaccione, Alberto Acquadro, Cinzia Comino, Cristiano Carli, Lorenzo Barchi, Matteo Martina

## Abstract

Globe artichoke (*Cynara cardunculus* var. *scolymus* L.) comprises a broad range of local ecotypes and varietal groups whose genetic diversity has been investigated through different molecular markers. However, recent advances in next-generation sequencing and pangenomics approaches provide new opportunities to capture genome-wide variation at higher resolution and to develop practical tools for varietal discrimination, traceability, and germplasm conservation. In this study, we developed the first pangenomic framework for cultivated artichoke and evaluated pangenome-informed SNP markers for varietal fingerprinting. Whole-genome resequencing data from the Italian local ecotype ‘Asti Sorì’ were integrated with publicly available genomic data from representative globe artichoke and cultivated cardoon accessions to construct and annotate a pangenome. Genome-wide SNP and presence/absence variation (PAV) analyses were combined with pangenome-anchored genotyping-by-sequencing (GBS) data from 45 accessions representing the main cultivated varietal groups. The pangenome revealed a largely conserved core gene repertoire alongside a smaller accessory component, with gene accumulation curves suggesting a tendency toward saturation within the sampled cultivated germplasm. SNP- and PAV-based analyses provided complementary views of accession relationships and consistently resolved the principal cultivated groups. Across the broader germplasm panel, pangenome-anchored GBS-derived SNPs identified well-supported phylogenetic clusters corresponding to recognized varietal types. A reduced panel of 50 SNPs, selected through iterative random subsampling, retained at least 90% of the genetic diversity captured by the full dataset and reproduced its main population structure. This compact pangenome-anchored marker set provides a practical foundation for varietal fingerprinting, DUS-oriented applications, traceability, and conservation of traditional globe artichoke germplasm. Validation across independent collections will be required before routine deployment.

## 1 INTRODUCTION

Pangenome approaches are reshaping how intraspecific genomic diversity is studied in crop species. By integrating sequence data from multiple accessions, pangenomes provide access to genomic content not represented in any single reference assembly - including structural variants, presence/absence variation (PAV), and accession-specific sequences (Bayer et al., 2020; Golicz et al., 2016; Schreiber et al., 2024). In crops characterized by complex domestication histories, clonal propagation, or pronounced agro-ecological differentiation, single-genome references are increasingly recognized as inadequate representations of species-wide diversity (Chapman et al., 2022; Schreiber et al., 2024). Pangenomic frameworks have been applied to improve marker discovery, characterize population structure, and identify loci associated with traits of agronomic importance across a range of vegetable and fruit crops (Li et al., 2023; Liu et al., 2023; Zhou et al., 2022).

Globe artichoke (*Cynara cardunculus* var. *scolymus* L.) is a diploid, cross-pollinating member of the Asteraceae family, primarily cultivated across the Mediterranean basin, with an estimated genome size of approximately 1.07 Gb (Acquadro et al., 2020). Its predominantly clonal propagation has contributed to the differentiation and maintenance of morphologically distinct varietal groups adapted to local agro-ecological conditions (Martina et al., 2024). Morpho-agronomic classifications identified major varietal groups based primarily on capitulum morphology and production traits, including “Catanese”, “Romanesco”, “Spinoso”, and “Violetti” types (Miccolis & Elia, 1996). Subsequent molecular studies based on SSRs, AFLPs, and genotyping-by-sequencing (GBS) approaches further refined this structure, distinguishing genetically differentiated groups broadly corresponding to “Catanese/Violetto di Provenza”, “Spinoso”, “Romanesco”, “Violetto di Toscana”, and “Macau” germplasm types (Pavan et al., 2018; Portis et al., 2025; Rau et al., 2022).

Local ecotypes occupy a distinct segment of crop diversity: maintained by farmers over generations through empirical selection in specific agro-ecological contexts, they often preserve allele combinations that are not present in improved or commercial varieties (Cheng et al., 2024). The globe artichoke ecotype ‘Asti Sorì’ has been cultivated in the Asti area of Piedmont (north-western Italy) for local fresh markets and is recognized by local producers for its distinctive agronomic characteristics. Despite its historical presence in the territory, ‘Asti Sorì’ has not been characterized at the genomic level, and no molecular tools are available for its formal identification or protection.

No pangenomic resource currently exists for globe artichoke, limiting the completeness of diversity analyses and the characterization of non-reference genomic content. At the same time, the absence of standardized SNP-based marker panels constrains varietal registration, DUS (distinctness, uniformity, and stability) testing, and the molecular traceability of traditional ecotypes. Previous genotyping-by-sequencing studies based on RADseq-derived markers demonstrated the potential of genome-wide SNP discovery for the analysis of globe artichoke population structure and diversity (Acquadro et al., 2016; Comino et al., 2016), but these approaches were not anchored to a pangenomic framework and were not specifically optimized for reduced diagnostic marker panel development. Accession-specific variants identified from limited resequencing panels may provide an initial route for marker discovery; however, loci appearing private within small discovery sets may not retain discriminatory power when evaluated across broader germplasm collections. A population-level approach based on pangenome-anchored GBS-derived SNPs therefore provides a more scalable route toward practical varietal fingerprinting tools.

In this study, we used the Italian local ecotype ‘Asti Sorì’ as a case study and genomic entry point to develop a first pangenomic framework for globe artichoke. Specifically, we: (i) generated whole-genome resequencing data for one representative ‘Asti Sorì’ accession and integrated it with publicly available resequencing data from representative *C. cardunculus* accessions, including cultivated globe artichoke types and one cultivated cardoon, together with the globe artichoke reference genome; (ii) reconstructed and annotated accession-level genomes using a reference-guided approach and used them to build a first pangenomic framework; (iii) characterized genome-wide SNP and presence/absence variation across the resequenced accessions, including accession-specific variants as preliminary candidate markers; and (iv) assessed pangenome-anchored GBS-derived SNPs in a broader globe artichoke germplasm panel to identify a reduced marker set for varietal fingerprinting.

## 2 MATERIALS AND METHODS

### 2.1 Plant material and agronomic evaluation

The local globe artichoke ecotype ‘Asti Sorì’ was evaluated under field conditions in two distinct environments in the Province of Asti (Piedmont, north-western Italy), characterized by different pedoclimatic conditions and cultivation practices. Experimental trials were conducted in the municipalities of Costigliole d’Asti and Revigliasco, including approximately 600 and 400 clonally propagated plants, respectively. At each site, ten representative plants were selected for detailed phenotypic assessment. Standard local agronomic practices were applied, including seasonal sucker removal to regulate vegetative growth and maintain plant productivity. Phenotypic observations included plant vigor, head number per plant, head weight, and head morphology (diameter, height, and shape), together with general observations on phenological development and plant health status throughout the growing season. The agronomic evaluation was performed to document the phenotypic consistency of the local material across environments and to support the genomic characterization of the ‘Asti Sorì’ ecotype used in subsequent analyses. To assess genetic variation within varietal types, a total of 45 globe artichoke accessions derived from a broader globe artichoke germplasm collection representative of the major cultivated varietal groups (Comino et al., 2016; Rau et al., 2022) were selected for DNA extraction and subsequent molecular analyses. Globe artichoke accessions were subdivided into seven groups based on morpho-phenotypic traits, primarily related to head characteristics (Supplementary Table S1).

### 2.2 DNA extraction, sequencing, and data collection

Genomic DNA was extracted from fresh/lyophilised young leaves of each accession using a modified CTAB protocol (Doyle, 1991; Li et al., 2024). RNAse A was used to remove RNA contamination, and DNA quality was checked by 1% (w/v) agarose gel electrophoresis. Quantification was performed by Qubit 2.0 (Life Technologies, Carlsbad, CA, USA), based on Qubit dsDNA BR Assay (Life Science), following Martina et al., 2023. Whole-genome sequencing of ‘Asti Sorì’ was performed at approximately 35× coverage using an Illumina NovaSeq X platform with 150PE reads insert size. In addition, publicly available resequencing data from representative Italian globe artichoke varietal types ‘Violetto di Sicilia’ (VS), ‘Violetto di Toscana’ (VT), ‘‘Romanesco C3’ (C3), ‘Spinoso di Palermo’ (SP), and the cultivated cardoon ‘Altilis 41’ (A41) were retrieved from the NCBI Sequence Read Archive (SRA - PRJNA238069; Acquadro et al., 2017). These materials included five cultivated globe artichoke accessions and one cultivated cardoon accession, and were used for genome reconstruction, comparative analyses, and pangenome construction.

### 2.3 Reference-guided genome reconstruction and annotation

Genome reconstruction of the selected varietal types was performed using a reference-guided approach. Cleaned sequencing reads were assembled de novo using MEGAHIT (Li et al., 2015) and resulting contigs were scaffolded against the globe artichoke reference genome (V2; Acquadro et al., 2020) using RagTag (Alonge et al., 2022). Contigs shorter than 500 bp were discarded. Structural gene annotation was carried out using a combined approach integrating ab initio prediction and transcript evidence. Gene models were predicted using Helixer and BRAKER3 (Gabriel et al., 2024; Holst et al., 2026; Stiehler et al., 2021), incorporating publicly available transcriptomic datasets (Scaglione et al., 2012). Functional annotation was performed using DIAMOND (Buchfink et al., 2021) against curated protein databases.

### 2.3 Pangenome construction

A pangenome was constructed by integrating the reconstructed genomes of all accessions with the reference genome. De novo assemblies from all reconstructed accessions were aligned to the reference using minimap2 (Li, 2021), and non-aligned sequences and unaligned regions longer than 500 bp were extracted as candidate novel sequences. Redundant sequences were collapsed using MMseqs2 (Steinegger and Söding, 2017), and contaminant sequences were filtered through similarity searches against public nucleotide databases. The final non-redundant sequences were merged with the reference genome to generate the globe artichoke pangenome. Pangenome expansion dynamics were modeled using Heaps’ law, and core genome size was estimated using an exponential decay model, as implemented in PanGP (Zhao et al., 2014). Genes were classified into core, softcore, shell, and cloud categories based on their frequency across accessions.

### 2.4 Functional annotation and GO enrichment analysis

Protein sequences were extracted from the pangenome annotation using gffread. Functional annotation was performed using four complementary approaches: (i) eggNOG-mapper (v2.1.13; Cantalapiedra et al., 2021) with the eggNOG 5.0 database, restricting orthology inference to Viridiplantae (taxid 33090); (ii) InterProScan (v5.67-99.0; Jones et al., 2014) against all integrated member databases; (iii) DIAMOND (v2.0.15; Buchfink et al., 2021) BLASTp against the UniProt Swiss-Prot reviewed protein database (downloaded April 2025, e-value ≤ 1×10⁻⁵); and (iv) DIAMOND BLASTp against the NCBI RefSeq plant protein database (e-value ≤ 1×10⁻⁵). Functional attributes - including GO terms, KEGG pathways, COG categories, and descriptive annotations - were merged by gene identifier into a unified functional table. GO terms were assigned with priority to InterProScan and eggNOG-derived terms, supplemented by Swiss-Prot hits. A species-specific OrgDb annotation package was built using the AnnotationForge package (v1.46.0) in R, incorporating gene-to-GO, gene-to-pathway, and gene-to-symbol mappings for all pangenome genes. GO enrichment analyses were performed in R using the clusterProfiler package (v4.12.0; Yu et al., 2012) with the enrichGO function (BH-adjusted p-value cutoff 0.05; q-value cutoff 0.20), using the full set of annotated pangenome genes as the statistical universe. Redundant GO terms were removed using the simplify function (similarity cutoff 0.7). For the ‘Asti Sorì’-unique gene set (n = 33), a relaxed threshold (BH p < 0.10) was applied given the limited gene number. To assess the functional stratification of the pangenome along the core-to-cloud axis, curated macro-categories of biological function were defined based on six anchor GO term sets (defense and immunity, specialized metabolism, signal transduction, transport, transcriptional regulation, primary metabolism and housekeeping), expanded to all descendant terms using GO ontology offspring relationships. Enrichment of each macro-category within each pangenome compartment (core, softcore, shell, cloud) relative to the annotated pangenome universe was assessed using Fisher’s exact test with BH correction.

### 2.5 Variant calling and population analysis

Sequencing reads from each accession were aligned to the pangenome using BWA-MEM2 (Vasimuddin et al., 2019). Variant calling was performed using bcftools (Danecek et al., 2021), applying standard quality filters (minimum mapping quality, read depth, and missing data thresholds). Genetic relationships among accessions based on SNP variation were investigated using principal component analysis (PCA) implemented in SNPRelate (Zheng et al., 2012) and maximum-likelihood phylogenetic inference using IQ-TREE2 (Minh et al., 2020). Presence/absence variation (PAV) analyses were performed using SGSGeneLoss (Golicz et al., 2015), and PAV-based phylogenetic relationships were compared with SNP-derived clustering patterns to evaluate complementary patterns of genome diversity across accessions.

### 2.6 GBS-based genotyping and minimum SNP panel for varietal fingerprinting

To extend comparative analyses to a broader diversity panel, a subset of 45 globe artichoke accessions was selected from a previously characterized germplasm collection representative of the major cultivated varietal groups (Comino et al., 2016; Rau et al., 2022). The selected accessions encompassed the seven principal morpho-agronomic typologies currently recognized within cultivated globe artichoke germplasm: “Catanesi” (CAT), “Violetti di Provenza” (VPR), “Violetti di Toscana” (VTO), “Romaneschi” (ROM), “Macau” (MAC), “Spinosi” (SPI), and “Green et al.” (GEA). Genotyping-by-sequencing was performed using a two-enzyme RADseq approach based on HindIII and NlaIII digestion, with minor modifications from the protocol described by Acquadro et al. (2016). Briefly, genomic DNA was digested with HindIII, ligated to barcoded P1 adapters, pooled, and subsequently digested with NlaIII before ligation to P2 Illumina indexed adapters. Libraries were purified, enriched through PCR to incorporate Illumina sequencing and indexing sites, and sequenced on an Illumina HiSeq 1000 platform according to the manufacturer’s protocol.

Sequencing reads were quality filtered and aligned to the globe artichoke pangenome generated in this study using BWA-MEM2 (Vasimuddin et al., 2019). SNP calling was performed using bcftools mpileup/call utilities (Danecek et al., 2021), applying standard quality and depth filters. Pangenome-anchored SNPs were used to investigate genetic relationships among accessions through principal component analysis and maximum-likelihood phylogenetic reconstruction using IQ-TREE2 (Minh et al., 2020). Population stratification within the broader diversity panel was further explored using sparse non-negative matrix factorization (sNMF) implemented in the LEA package (Frichot and François, 2015), evaluating K values from 1 to 20 with 20 independent runs per K.

A random SNP subsampling approach was then used to identify reduced marker panels representative of the genetic diversity captured by the full GBS dataset. Starting from the filtered biallelic SNP VCF file, genotype calls were processed in R using the *vcfR* package (Knaus and Grünwald, 2016) and converted into numeric allele dosage values, with homozygous reference, heterozygous, and homozygous alternative genotypes coded as 0, 1, and 2, respectively. Missing genotypes were retained. A reference pairwise genetic distance matrix was first calculated from the complete SNP dataset using dosage-based Euclidean distances; for each accession pair, distances were computed only on markers with non-missing genotypes in both accessions and rescaled to account for the number of available markers. For each target panel size, corresponding to 50, 75, and 100 SNPs, 100 random SNP subsets were generated without replacement. For each subset, the same distance metric was recalculated and compared with the full-dataset distance matrix using Pearson correlation across all accession pairs. For each panel size, the subset showing the highest correlation with the full-dataset distance matrix was retained as the best representative reduced marker panel. The selected SNPs were reported as genomic coordinates and extracted from the original VCF file using *bcftools* (Danecek et al., 2021) for downstream phylogenetic reconstruction and fingerprinting assessment.

## 3 RESULTS

### 3.1 Phenotypic stability of the ‘Asti Sorì’ ecotype across environments

The local globe artichoke ecotype ‘Asti Sorì’ is characterized by green, elongated, and spineless capitula, representative of a traditional Northern Italian varietal type (Figure 1).

**Figure 1.**
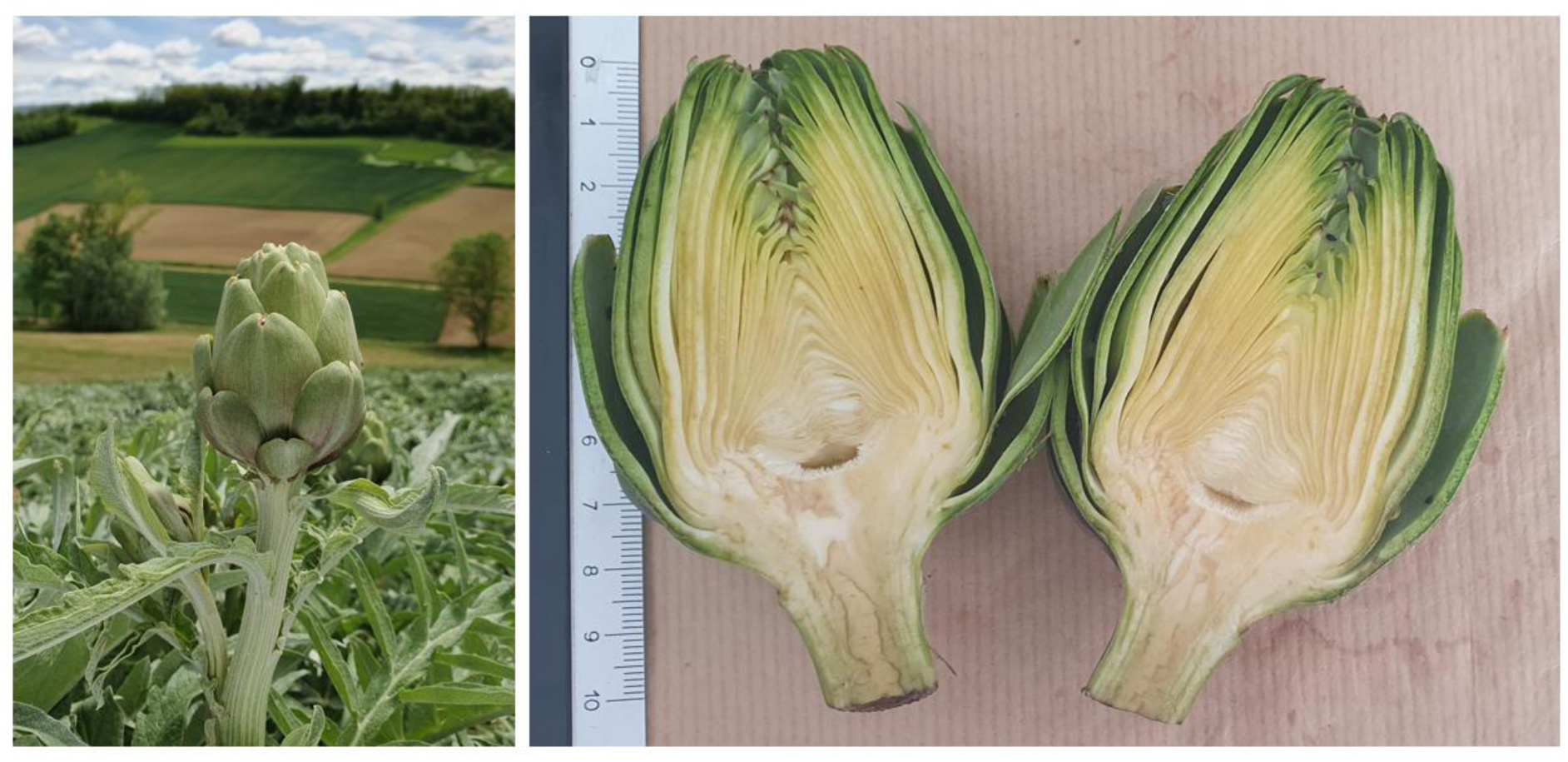
Representative morphology of the globe artichoke ecotype ‘Asti Sorì’. Left: field-grown plant showing the characteristic green, elongated, and spineless capitulum cultivated in the Asti area (Piedmont, north-western Italy). Right: longitudinal section of a representative capitulum highlighting the morphology of the receptacle and inner bracts.

Agronomic evaluation across two contrasting environments in the Asti area revealed limited phenotypic variation, supporting the stability of the material under different pedoclimatic conditions. Plants cultivated in Costigliole d’Asti showed greater variability in vegetative traits, particularly plant height and stem diameter, than those grown in Revigliasco (Figure 2). In contrast, yield-related traits displayed overlapping distributions between environments. The most consistent pattern was observed for capitulum morphology traits, including head dimensions and shape index (L/R ratio), which showed limited variation and narrow trait distributions across sites. Taken together, these observations document the agronomic consistency of the ‘Asti Sorì’ material across contrasting environments. The limited phenotypic variation observed is consistent with clonal propagation and supports the suitability of this local ecotype as a defined target for genomic characterization and molecular traceability.

**Figure 2.**
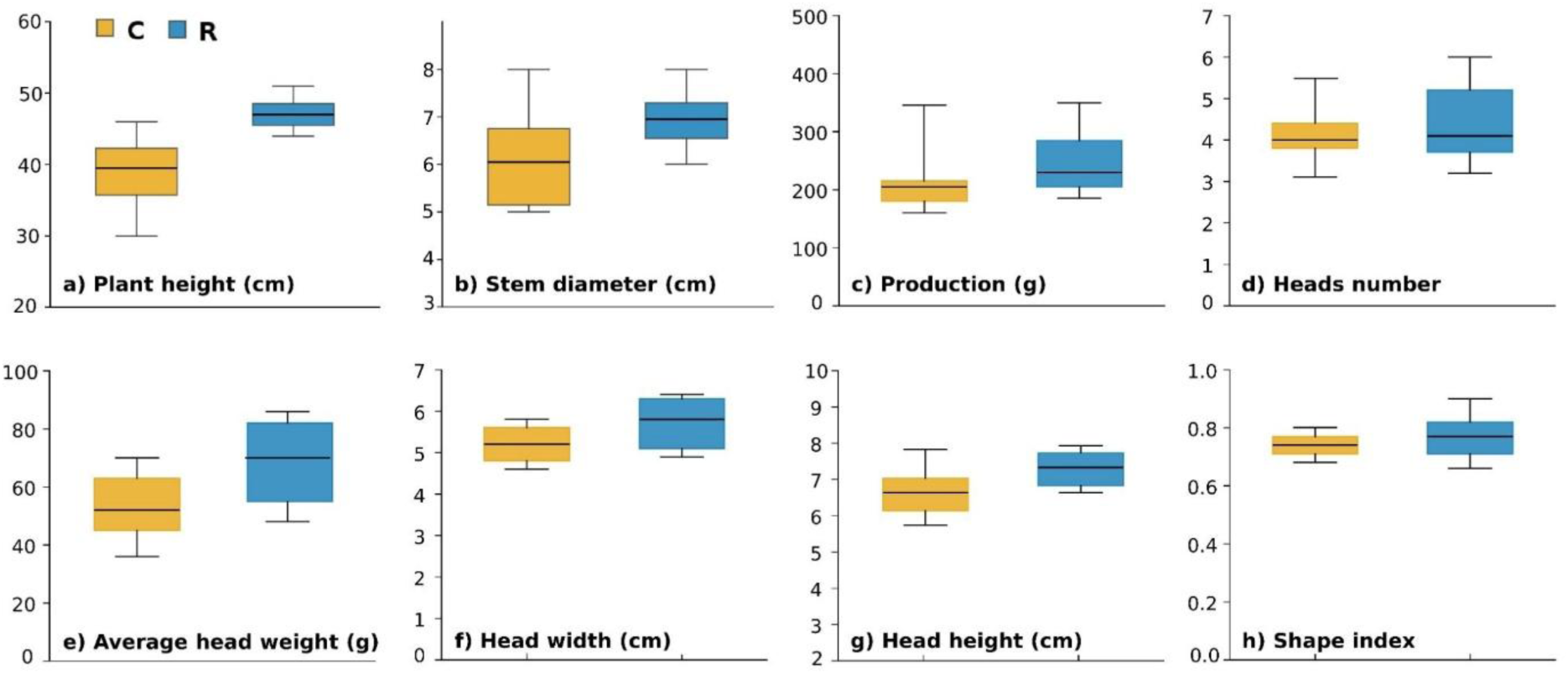
Phenotypic traits of the ‘Asti Sorì’ ecotype evaluated in the two experimental environments, Costigliole d’Asti (C, orange) and Revigliasco (R, blue). (a) plant height; (b) stem diameter; (c) total production per plant; (d) number of capitula per plant; (e) average head weight; (f) capitulum width; (g) capitulum height; and (h) capitulum shape index (L/R ratio).

### 3.2 Reference-guided genome reconstruction and annotation

A reference-guided assembly approach enabled the generation of reconstructed genomes for all accessions, including the newly sequenced ‘Asti Sorì’, which were subsequently annotated using a combination of ab initio prediction and transcript-supported methods. The resulting assemblies were produced after the RagTag reconstruction and annotated using a hybrid approach by combining machine learning approach and transcriptomic data. Structural annotation allowed the identification of the reconstructed genomes, identifying between a minimum of 34,074 (VT) and a maximum of 34,746 (C3) genes (Table 1). The combined use of Helixer and BRAKER3, supported by transcriptomic data (Scaglione et al., 2012), resulted in an expanded gene set compared to previous annotations (Acquadro et al., 2020). Overall, these reconstructed and annotated genomes provided the basis for subsequent comparative analyses and pangenome construction.

**Table 1.**
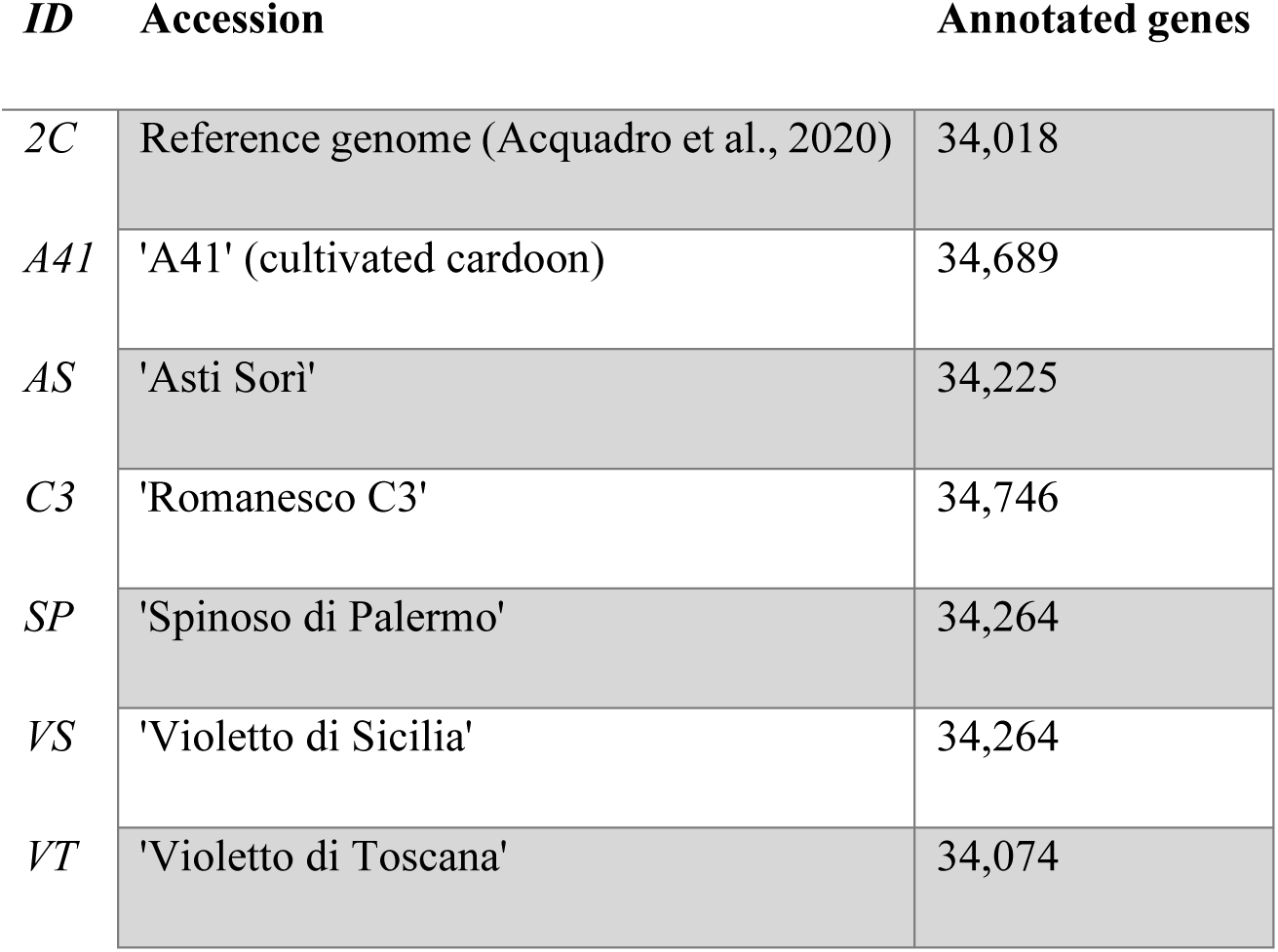
Number of annotated genes per accession genome included in the globe artichoke pangenome (Helixer + BRAKER3).

### 3.3 Pangenome construction and architecture

To capture genome-wide diversity in the analyzed *C. cardunculus* material, a pangenome was constructed by integrating the reconstructed genomes of five globe artichoke accessions and one cultivated cardoon accession with the globe artichoke reference genome. Following redundancy removal and contamination filtering, a total of 131.3 Mb of non-reference sequences was identified, contributing additional genomic content to the artichoke pangenome. These sequences expanded the genomic space beyond the reference assembly, highlighting the presence of previously uncharacterized regions across accessions. Gene prediction on non-reference sequences identified 2,894 novel protein-coding genes. Among these, 1,352 were supported by conserved domains (InterPro), and 1,023 were assigned functional annotations including Gene Ontology (GO) terms. The integration of these genes with those of the reference genome resulted in a pangenome comprising a total of 37,198 protein-coding genes and a genome size of approximately 0.86 Gb.

Based on gene presence across accessions, the pangenome was partitioned into core, softcore, shell, and cloud gene categories (Figure 3). A total of 30,817 genes were classified as core, being shared by all accessions, 1,874 as softcore, shared by six accessions, 3,471 as shell, shared by two to five accessions, and 580 as cloud genes, present in only one accession (Supplementary Table S2). The complete list of accession-specific cloud genes and their functional annotation is provided in Supplementary Table S3. Most genes were classified as core or softcore, indicating a high level of conservation across the analyzed germplasm, while a smaller fraction constituted the accessory genome, including shell and cloud components. Among the analyzed accessions, ‘A41’ showed the highest proportion of cloud genes (0.70%), whereas ‘VT’ exhibited the lowest (0.07%; Table 2; Supplementary Table S2; Figure 3).

**Table 2.** Percentage of genes assigned to each pangenome category (core, softcore, shell, cloud) per accession.

| Class | 2C (%) | A41 (%) | AS (%) | C3 (%) | SP (%) | VS (%) | VT (%) |
| --- | --- | --- | --- | --- | --- | --- | --- |
| Core | 90.59 | 88.84 | 90.04 | 88.69 | 89.94 | 89.94 | 90.44 |
| Soft | 3.62 | 4.47 | 5.07 | 5.13 | 4.88 | 4.78 | 4.79 |
| Shell | 5.42 | 5.99 | 4.79 | 6.02 | 5.07 | 5.12 | 4.69 |
| Cloud | 0.37 | 0.70 | 0.10 | 0.16 | 0.12 | 0.17 | 0.07 |

**Figure 3.**
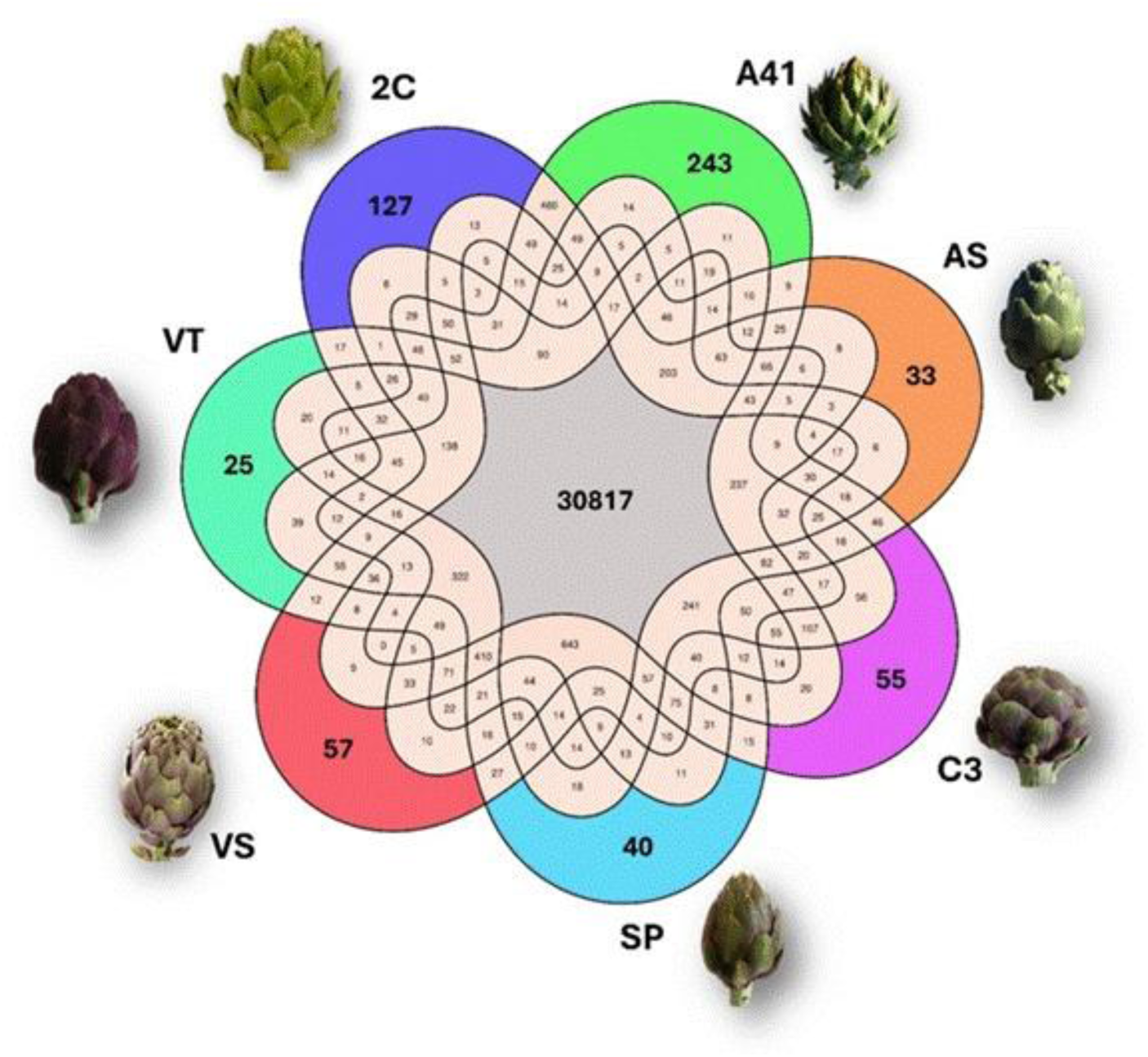
Gene presence/absence relationships among the seven genomes included in the globe artichoke pangenome. Numbers within the colored sectors indicate accession-specific (cloud) genes, whereas overlapping regions correspond to genes shared by different combinations of accessions. Accessions included in the analyses were: ‘Asti Sorì’ (AS), ‘Romanesco C3’ (C3), ‘Violetto di Toscana’ (VT), ‘Violetto di Sicilia’ (VS), ‘Spinoso di Palermo’ (SP), cultivated cardoon ‘Altilis 41’ (A41), and the globe artichoke reference genotype ‘2C’.

The dynamics of pangenome expansion were assessed by modeling gene accumulation curves (Figure 4). The pangenome size increased with the addition of new genomes, reaching 37,198 genes with the inclusion of all seven genomes. The core genome, representing genes shared by all accessions, was estimated at 30,817 genes and decreased as additional genomes were incorporated, consistent with the expected behavior under Heap’s law. Although the rate of new gene discovery decreased with each additional genome, indicating a tendency toward saturation within the sampled cultivated germplasm, the limited number of genomes included prevents definitive conclusions about the pangenome status of *C. cardunculus* across its full taxonomic and geographic range.

**Figure 4.**
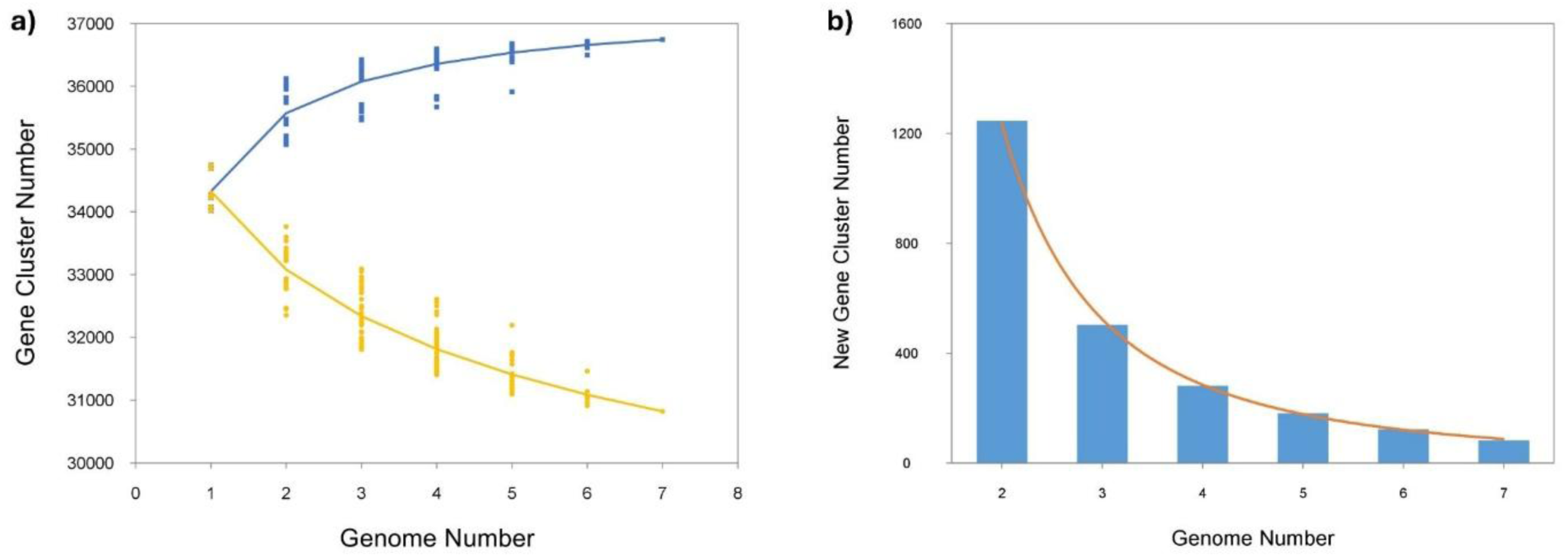
(a) Gene accumulation curves of the pan-genome (blue) and the core genome (orange). These curves describe the number of new genes (pan-genome) and genes in common (core genome) obtained by adding a new genome to a previous set. The blue vertical lines denote the pan-genome size for each genome for comparison. The orange vertical lines show the core genome size for each genome for comparison. The curve is the least squares fit of the power law for the average values. (b) Curve (orange) for the number of new genes at each increase in the number of genomes.

Overall, the pangenome expansion curve showed a tendency toward saturation within the sampled cultivated germplasm, with a decreasing rate of novel gene discovery as additional accessions were included. This pattern consisted of a conserved gene repertoire among the analyzed accessions, accompanied by a limited accessory component contributing to gene-content variation.

To characterize the functional landscape of the globe artichoke pangenome, GO enrichment analysis was performed for each gene category using a custom Cynara cardunculus annotation database built from eggNOG-mapper, InterProScan, and DIAMOND-based similarity searches against Swiss-Prot and NCBI RefSeq plant protein databases (see Materials and Methods). The pangenome displayed a marked functional stratification along the core-to-cloud axis (Figure 5). Transcriptional regulators and transporter-encoding genes were significantly enriched in the core genome (BH-corrected Fisher’s test, p < 0.001), consistent with a conserved regulatory and metabolic backbone shared across all accessions. In contrast, genes involved in specialized (secondary) metabolism and defense responses showed a monotonically increasing enrichment from core to cloud categories, with the strongest signal observed for specialized metabolism (shell: p < 0.001; cloud: p < 0.01). Genes assigned to primary metabolism and housekeeping functions were uniformly distributed across all pangenome categories (p > 0.05), confirming that the observed gradient reflects genuine functional partitioning rather than annotation coverage biases.

**Figure 5.**
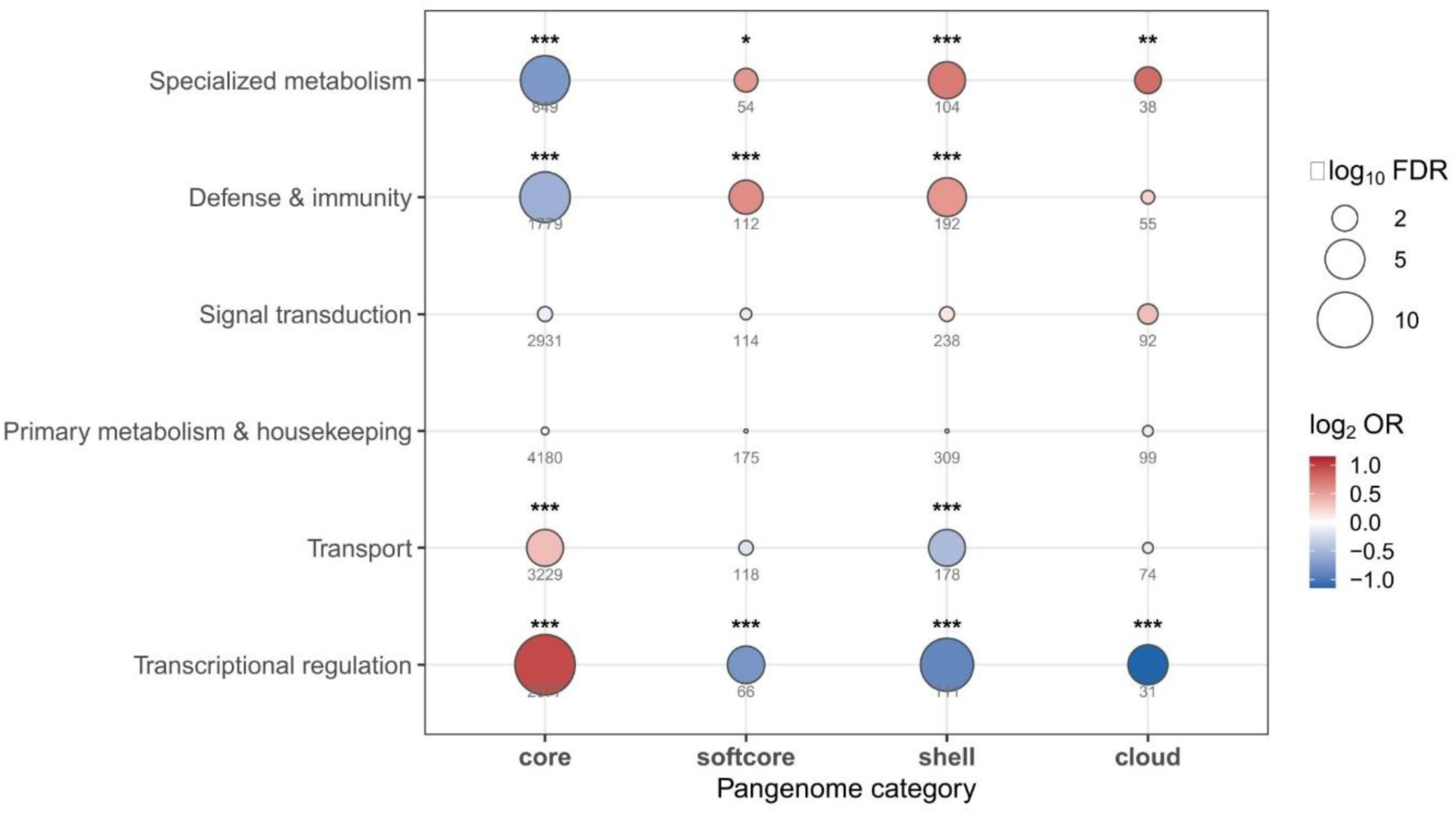
Functional stratification of the globe artichoke pangenome. Enrichment of curated functional categories along the core-to-cloud axis. Bubble size is proportional to enrichment significance (−log₁₀ FDR), color indicates the enrichment odds ratio (log₂ OR), and numbers correspond to the number of enriched genes. Asterisks indicate statistically significant enrichments.

Detailed GO enrichment analysis of the accessory genome (shell and cloud genes combined; n = 4,051) further resolved the biological functions associated with accession-variable genes (Figure 5; Supplementary Table S4). Within biological processes, isoprenoid and terpenoid biosynthetic processes were among the most significantly enriched terms (BH p < 0.01), consistent with the central role of sesquiterpene lactones and phenylpropanoids in globe artichoke quality traits and inter-accession chemical diversity. Nitrate assimilation and nitric oxide biosynthesis were also enriched, pointing to variation in nitrogen metabolism and signaling across accessions. Among molecular function terms, transmembrane receptor tyrosine kinase activity and calcium channel activity showed the strongest enrichment, reflecting the diversity of immune receptor and signal transduction gene repertoires in the variable portion of the pangenome. Cellular component analysis revealed enrichment of photosynthetic membrane and plastid ribosomal subunit terms, indicating structural variation in plastid-associated gene content (Supplementary Figure 1).

Accession-specific functional signatures were explored for the ‘Asti Sorì’ (AS) and the cultivated cardoon ‘A41’ accessions. Among the 33 genes uniquely present in ‘Asti Sorì’, GO enrichment analysis at a relaxed significance threshold (FDR < 0.10) identified terms associated with specialized metabolism, including sesquiterpene synthase activity (GO:0010334) and epoxide hydrolase activity, suggesting a distinctive secondary metabolite capacity in this ecotype. However, the limited number of accession-unique genes and the absence of significant enrichment at the conventional FDR < 0.05 threshold preclude firm conclusions. For the cultivated cardoon ‘A41’, functional annotation coverage was substantially reduced: 83% of the 243 A41-unique genes corresponded to non-reference pangenome gene models characterized by short, predicted protein sequences (median length 105 amino acids, compared with 310 amino acids for reference-derived genes), a limitation inherent to de novo assembly-based prediction. Only a single GO term reached significance for A41-unique genes (oxidoreduction-driven active transmembrane transporter activity; FDR = 0.004), precluding broader functional inference for the cardoon-specific gene set.

### 3.4 Genome-wide variation, accession relationships, and candidate diagnostic markers

Genome-wide variant calling revealed extensive genetic variation across the analyzed accessions, with a predominance of heterozygous variants, consistent with the allogamous nature of *C. cardunculus*. The total number of variants differed among accessions, with the cultivated cardoon accession ‘A41’ showing the lowest number of variants (2.9 million) and the ‘Asti Sorì’ accession the highest (9.7 million - Figure 6).

**Figure 6.**
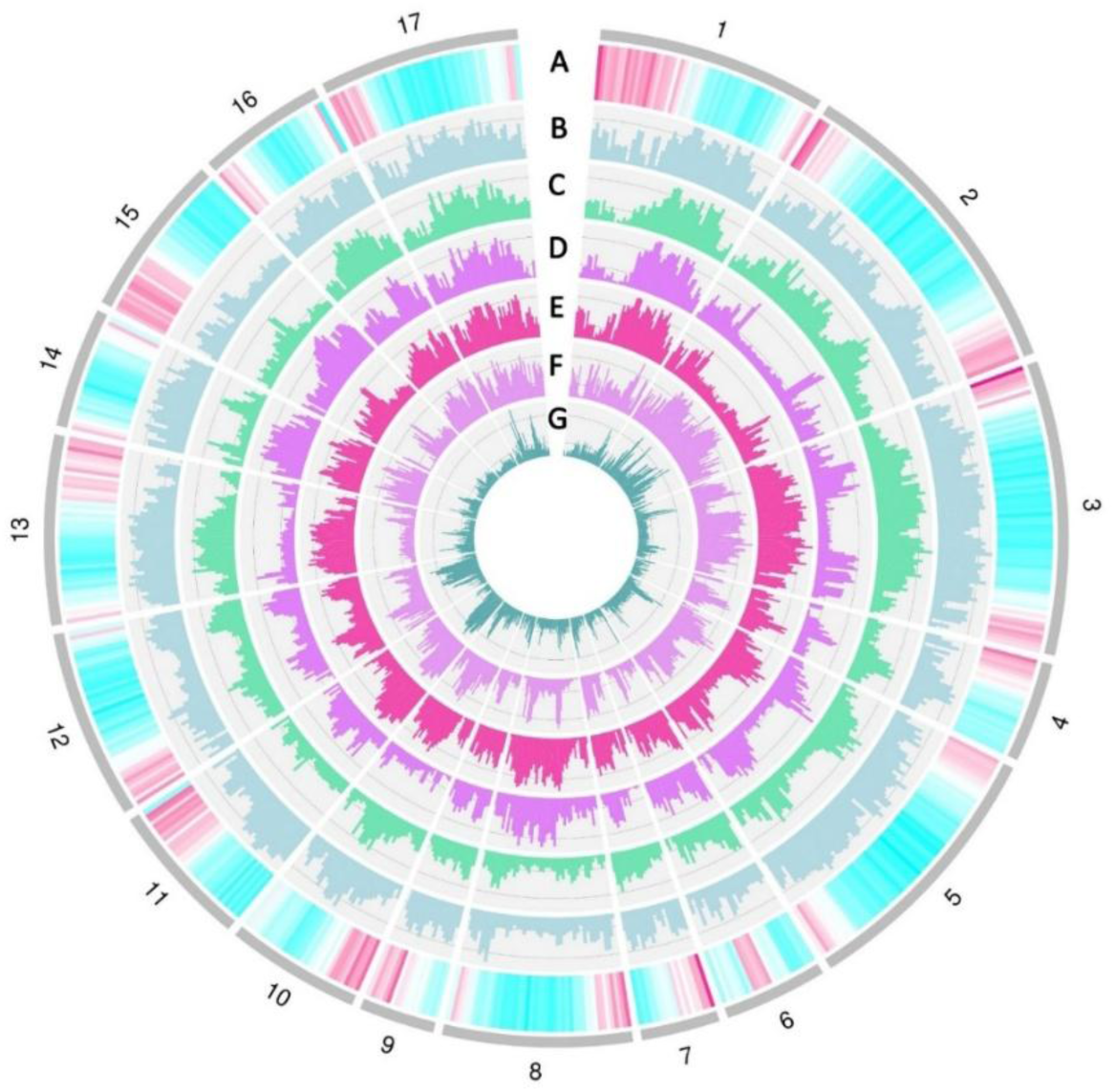
Circus diagram showing gene density in the reference genome (A) and the density of heterozygous SNPs in the ‘Asti Sorì’ artichoke (B), ‘Romanesco C3’ (C), ‘Violetto di Toscana’ (D), ‘Violetto di Sicilia’ (E), ‘Spinoso di Palermo’ (F), and the cultivated cardoon ‘A41’ (G).

Genetic relationships among accessions were investigated using both SNP-based and presence/absence variation (PAV)-based approaches derived from the pangenome. Principal component analysis (PCA) based on genome-wide SNPs revealed a clear genetic differentiation among accessions (Supplementary Figure 2A). The first two principal components explained a substantial proportion of the total variance (29.35% and 17.22%, respectively). The reference accession ‘2C’ and the cultivated cardoon ‘A41’ were clearly separated along the first principal component, indicating distinct genetic profiles. In contrast, ‘VT’, ‘VS’, and ‘SP’ clustered closely together, suggesting a high degree of genetic similarity. The ‘Asti Sorì’ accession occupied a distinct intermediate position, whereas ‘C3’ showed greater divergence along the second component.

Maximum-likelihood phylogenetic inference based on 34,583 pruned SNPs supported the clustering pattern observed in the PCA (Supplementary Figure 2B). ‘A41’ and ‘2C’ formed a distinct clade, while ‘SP’, ‘VS’, and ‘VT’ grouped together with high bootstrap support. ‘Asti Sorì’ and ‘C3’ clustered separately and similar overall topology was observed using PAV-based phylogenetic reconstruction, although differences in the relative positioning of some accessions were detected.

Taken together, these analyses indicate that both nucleotide variation and gene presence/absence contribute to the differentiation of globe artichoke accessions. While SNP-based analyses primarily captured overall genetic similarity, PAV-based relationships highlighted complementary patterns of genome diversity associated with structural variation.

### 3.5 Varietal relationships and reduced SNP fingerprinting using pangenome-anchored GBS markers

Using the globe artichoke pangenome as a common coordinate framework, pangenome-anchored GBS-derived SNPs were analyzed across a panel of 45 accessions representative of the principal cultivated varietal groups. Principal component analysis based on the complete filtered SNP dataset revealed a clear genetic differentiation among varietal groups (Figure 7). The first principal component separated three major groups of accessions, with “Spinosi” materials positioned on the positive side of the axis, “Catanesi” and “Violetti di Provenza” accessions on the opposite side, and “Romanesco”, “Violetto di Toscana”, and related materials occupying intermediate positions. The second principal component further differentiated “Romanesco” and “Violetto di Toscana” accessions. The ‘Asti Sorì’ accession occupied a distinct position relative to all other groups, consistent with its Northern Italian origin and genomic distinctiveness.

**Figure 7.**
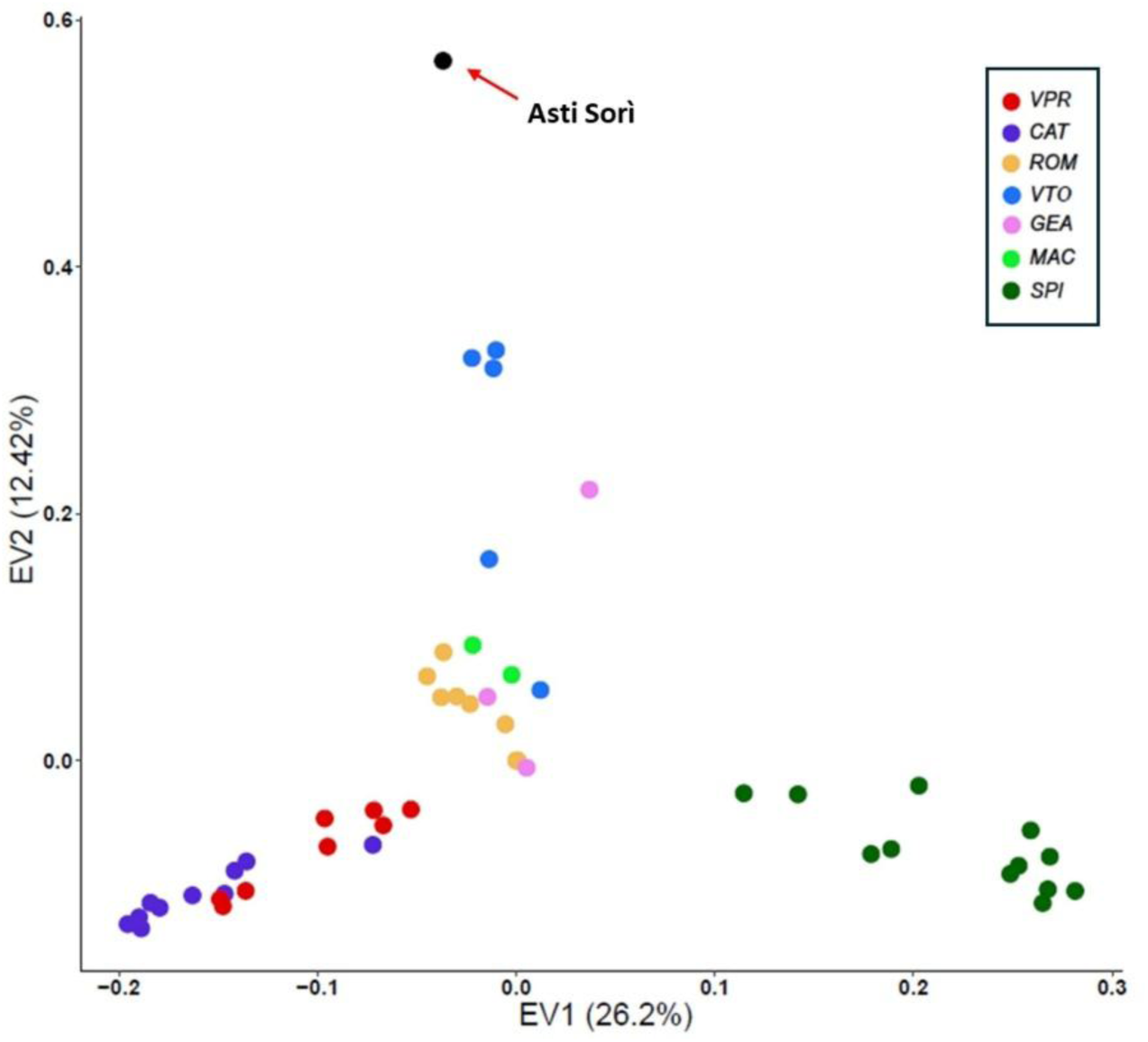
Principal component analysis (PCA) of the globe artichoke diversity panel based on the complete filtered set of pangenome-anchored GBS-derived SNPs. Accessions are colored according to the principal morpho-agronomic typologies as defined by Comino et al. (2016) and Rau et al. (2022): “Catanesi” (CAT), “Violetti di Provenza” (VPR), “Violetti di Toscana” (VTO), “Romaneschi” (ROM), “Spinosi” (SPI), “Macau” (MAC) and “Green et al.” (GEA). The “Asti Sorì” accession is indicated separately.

Population structure analysis resolved the analyzed accessions into distinct genetic groups broadly corresponding to the recognized morpho-agronomic typologies of cultivated globe artichoke (Figure 8, right side). Bayesian clustering analysis identified K = 6 as the most informative subdivision level, consistent with the complex structure previously reported for globe artichoke germplasm. “Catanesi” and “Violetti di Provenza” accessions clustered together with high assignment probabilities, confirming their close genetic relationship. Most “Romanesco” accessions and all “Violetto di Toscana” accessions formed clearly differentiated groups, whereas “Macau”, “Green et al.” (GEA), and two “Romanesco” accessions displayed more admixed genetic backgrounds, occupying intermediate positions among the principal clusters. “Spinosi” accessions formed a distinct genetic group that was further subdivided into two major subclusters corresponding to their geographic origin, separating Sicilian spiny accessions from the Sardinian ones. The ‘Asti Sorì’ accession showed a distinct and independent clustering profile relative to all other varietal groups, supporting its genomic uniqueness within the analyzed germplasm.

**Figure 8.**
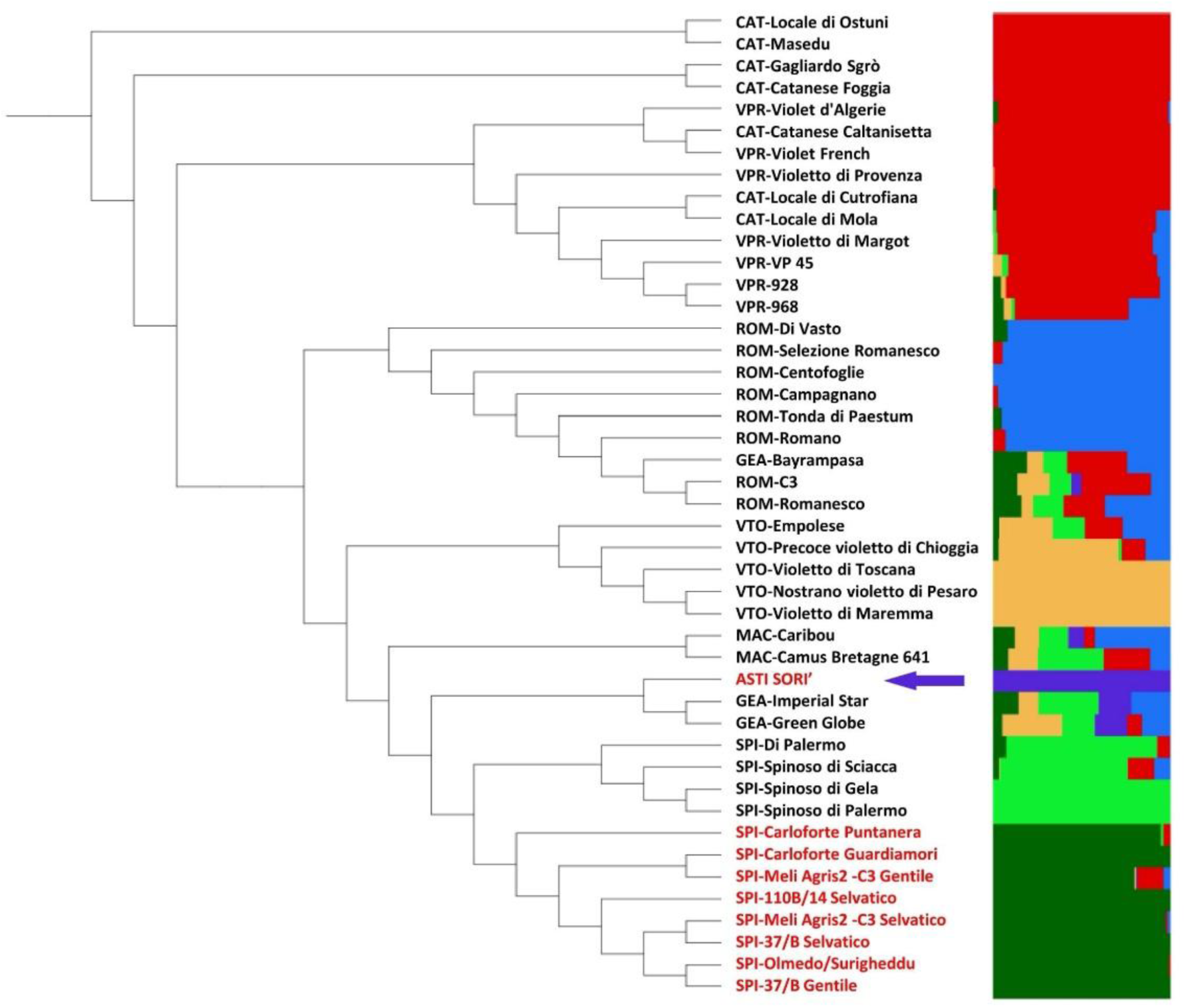
Population structure of the globe artichoke diversity panel based on pangenome-anchored GBS-derived SNPs. Right: Bayesian clustering analysis showing the assignment probabilities of individual accessions at K = 6. Left: maximum-likelihood phylogenetic tree inferred from the complete filtered SNP dataset. Accessions are labeled according to the principal morpho-agronomic typologies as in Fig. 7. Sardinian accessions are highlighted in red within the SPI group, “Asti Sorì” accession is indicated separately.

Maximum-likelihood phylogenetic reconstruction broadly supported the clustering patterns observed in both the PCA and Bayesian population structure analyses (Figure 8, left side). “Catanesi” and “Violetti di Provenza” accessions consistently clustered together while maintaining detectable internal differentiation.

“Romanesco” accessions formed a coherent phylogenetic group, including the two accessions that displayed partially admixed ancestry profiles in the Structure analysis. “Spinosi” accessions clustered into a distinct and well-supported clade, within which the subdivision between Sicilian and Sardinian materials was maintained, confirming the geographic structure previously highlighted by Bayesian clustering analyses. The ‘Asti Sorì’ accession grouped within the broader cluster of green-headed accessions, consistent with its morphological characterization as a green, spineless varietal type, while maintaining a distinct phylogenetic position relative to the remaining green-headed materials.

To evaluate the feasibility of reduced SNP-based varietal fingerprinting, iterative random subsampling analyses were performed on the complete pangenome-anchored SNP dataset. For panel sizes of 50, 75, and 100 SNPs, 100 random replicates were evaluated by Pearson correlation between the pairwise genetic distance matrix derived from each candidate panel and that obtained from the complete SNP dataset (Supplementary Table S6). The best-performing 50-SNP panel (Pearson’s r = 0.96; Supplementary Table S5) was retained for downstream validation. Phylogenetic reconstruction based exclusively on this 50-SNP subset largely recapitulated the principal varietal structure observed with the full dataset (Figure 9). Although some internal relationships among closely related accessions differed from those inferred using the complete SNP matrix, all major varietal groups remained recognizable, and each accession retained a distinct genetic profile. In particular, the ‘Asti Sorì’ accession maintained a clearly differentiated phylogenetic position, supporting the effectiveness of the reduced marker set for varietal discrimination and molecular fingerprinting within the analyzed germplasm.

**Figure 9.**
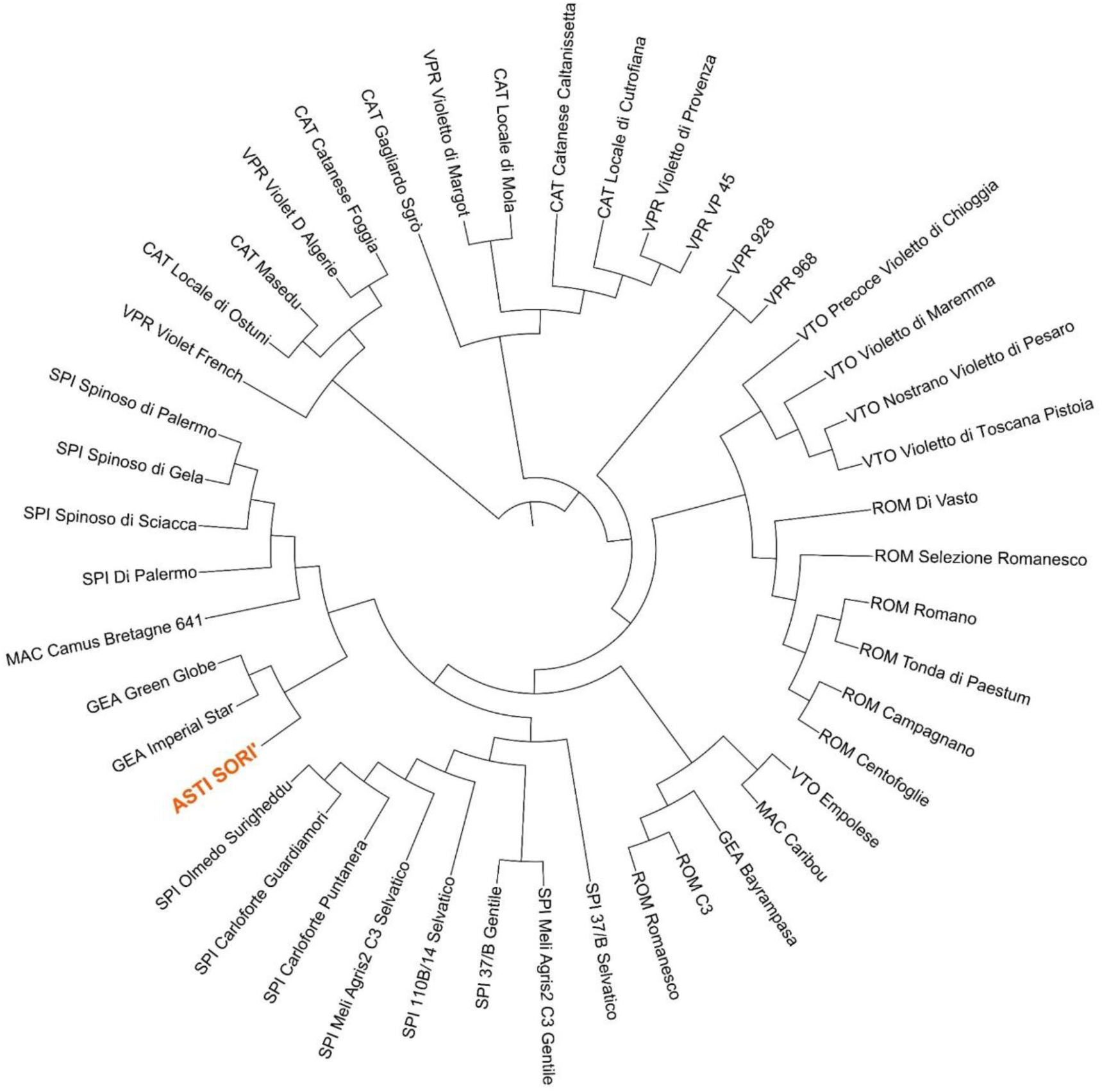
Radial phylogenetic tree inferred from the reduced set of 50 pangenome-anchored SNPs selected for varietal fingerprinting. Accessions are labeled according to the principal morpho-agronomic typologies reported in Figure 7, “Asti Sorì” accession is indicated in red.

## 4 DISCUSSION

We developed the first pangenomic framework for globe artichoke and explored SNP-based approaches for varietal discrimination. The globe artichoke pangenome revealed a genomic architecture dominated by a large, conserved core gene set, complemented by a smaller accessory component comprising genes variably distributed across accessions. This structure is broadly consistent with pangenomes reported in other crop species, where core gene content generally represents conserved developmental and metabolic functions, whereas accessory genes are often enriched for functions related to environmental response, secondary metabolism, disease resistance, and transposable element activity (Bayer et al., 2020; Bozan et al., 2023; Gaccione et al., 2025; Golicz et al., 2015). The pangenome accumulation curves suggested a tendency toward saturation within the sampled cultivated germplasm. However, the limited number of genomes analysed, all of cultivated origin, prevents strong conclusions about the overall openness of the *C. cardunculus* pangenome across its full taxonomic and geographic range. Inclusion of wild cardoon (*C. cardunculus* var. *sylvestris*) and more cultivated cardoon (*C. cardunculus* var. *altilis*) accessions in future pangenome projects will be important to capture the full extent of structural and sequence variation within this species complex. This first pangenomic framework therefore provides not only a genomic resource for globe artichoke, but also a baseline for distinguishing conserved species-level features from accession-variable genomic content.

The biological relevance of this pangenomic resource became evident when gene-content variation was interpreted in functional terms, revealing a clear functional stratification across pangenome categories. Transcriptional regulators and transporter genes were consistently enriched in the core genome, suggesting that conserved regulatory and transport-related functions represent part of the shared genomic infrastructure across the analysed accessions. In contrast, the accessory genome showed enrichment for specialized metabolism, particularly isoprenoid and terpenoid biosynthetic processes, together with defense-related functions including receptor kinase signaling and calcium channel activity. This distribution indicates that genes variably present across accessions are not merely residual or annotation-derived elements, but include functional categories potentially associated with adaptive and quality-related traits. A particularly relevant result was the increasing enrichment of specialized metabolism from softcore to cloud categories. This trend suggests that genes located at the periphery of the pangenome may preferentially contribute to accession-specific chemical diversity. Similar patterns have been reported in other crop pangenomes, including tomato and maize, where dispensable genome content is disproportionately associated with adaptive traits, disease resistance, and secondary metabolite diversification (Gao et al., 2019; Hirsch et al., 2014; Shi et al., 2026; Yang et al., 2025; Zhou et al., 2022). In globe artichoke, this observation is especially relevant because several secondary metabolites, including sesquiterpene lactones such as cynaropicrin and phenolic compounds such as chlorogenic acids, are central to product quality, nutritional value, and cultivar differentiation. The enrichment of terpenoid- and phenylpropanoid-related functions in the accessory genome therefore suggests that comparative pangenomics may provide a useful route to identify candidate genes contributing to chemical and adaptive diversity in *C. cardunculus*.

Population structure analyses based on genome-wide SNPs and presence/absence variation revealed broadly consistent, but not fully overlapping, patterns of accession relationships. This partial divergence is expected because SNP-based analyses capture nucleotide-level variation distributed across the genome, whereas PAV-based analyses describe differences in gene content. These two layers therefore provide complementary rather than redundant views of diversity, as also observed in other crops where structural and nucleotide variation can reflect partially independent evolutionary histories (Jayakodi et al., 2024; Qin et al., 2021; Zhou et al., 2022). Both approaches nevertheless consistently identified ‘Asti Sorì’ as distinct from the other resequenced accessions, reinforcing its genomic individuality within the analyzed material.

The high heterozygosity observed across the resequenced accessions, and particularly in ‘Asti Sorì’, is consistent with the allogamous reproductive biology of globe artichoke and with the complex allelic landscape expected in clonally maintained, outbreeding varieties. In such crops, vegetative propagation can maintain highly heterozygous genotypes over time, while local selection and farmer-mediated clonal maintenance may preserve combinations of alleles associated with agronomic performance or local adaptation. From this perspective, the genomic distinctiveness of ‘Asti Sorì’ should not be interpreted as evidence that the analysed accession exhaustively represents the ecotype, but rather as evidence that the local material contains a recognizable genomic component that can be targeted for further characterization.

These observations raised a practical question: whether the genomic variation detected in the pangenome could be translated into robust varietal discrimination tools. The broader GBS panel used in this study used a globe artichoke germplasm collection representing the major cultivated varietal groups (Comino et al., 2016; Rau et al., 2022). This resource provides an appropriate population-level context in which to test whether variants identified from resequencing data can contribute to varietal discrimination. The identification of accession-specific variants from the resequencing panel represented a first route toward candidate diagnostic markers for the other analysed accessions as well as for ‘Asti Sorì’. However, variants that appear private within a small discovery panel may not retain diagnostic value when broader germplasm diversity is considered. This is a general limitation of accession-specific marker discovery based on restricted genome panels and highlights the need to evaluate marker informativeness at the population scale.

For this reason, the study moved from private-variant discovery to a population-validated SNP fingerprinting strategy using the 45-accession GBS dataset. Genome-wide GBS-derived SNPs resolved the main varietal groups with high confidence, and a reduced set of 50 markers was sufficient to recapitulate the major axes of genetic differentiation observed in the full dataset. This result indicates that, within the analysed germplasm, a compact set of pangenome-anchored SNPs can capture the principal structure of cultivated globe artichoke diversity. The 50-SNP panel identified here therefore provides a computationally validated candidate resource for varietal fingerprinting in globe artichoke. The potential applications of this candidate marker set include support for DUS testing, product traceability, and conservation monitoring of traditional ecotypes. In DUS-oriented contexts, standardized molecular profiles may complement morphological descriptors, particularly for clonally propagated crops in which phenotypic evaluation can be influenced by environment and management conditions (Yang et al., 2021). For traceability, low-cost genotyping assays could support the verification of varietal origin along production chains. For conservation, molecular fingerprinting may help monitor the identity and stability of local ecotypes such as ‘Asti Sorì’, for which standardized genomic tools are currently lacking. In this sense, the integration of pangenome information with reduced SNP panels provides a practical framework for translating comparative genomics into applied varietal identification.

However, the 50-SNP panel should be considered a candidate resource rather than a ready-to-use diagnostic tool. Conversion of the selected SNPs into deployable genotyping assays, such as KASP probes or targeted amplicon sequencing, will require additional technical development and validation. More importantly, the discriminatory performance of the panel should be evaluated on independent germplasm collections before routine diagnostic use can be proposed. The 45-accession GBS panel analysed here, while representative of major Italian varietal groups, does not capture the full diversity of globe artichoke preserved in national and international germplasm repositories. In particular, the ability of the marker set to resolve closely related accessions within the same varietal group, a critical requirement for DUS-oriented applications, remains to be established.

Local ecotypes of crop species may harbor allele combinations shaped by long-term farmer selection under specific environmental and agronomic contexts, and their genomic characterization represents an important step toward molecular traceability, conservation, and formal recognition (Lazaridi et al., 2024; Ramirez-Villegas et al., 2022). In the case of ‘Asti Sorì’, field evaluation across two cultivation sites in the Asti area supported the agronomic consistency of the local material. This consistency is compatible with the clonal propagation of globe artichoke and supports the relevance of the ecotype as a defined target for molecular characterization, while not implying that a single accession fully captures its intra-ecotype diversity. Although only one representative ‘Asti Sorì’ accession was used for whole-genome resequencing and pangenome construction, this accession should be interpreted as a genomic entry point into the ecotype rather than as a complete description of its genetic breadth. Notably, ‘Asti Sorì’ was consistently resolved as a distinct accession in both WGS-based and GBS-based population analyses, supporting its individuality within the broader landscape of Italian globe artichoke diversity. In this framework, the fingerprinting analysis shows that ‘Asti Sorì’ is distinguishable from the other analysed materials, the pangenome indicates the presence of accession-specific gene-content features, while the reduced SNP panel provides a concrete route to monitor, protect, and valorise this local ecotype. These results highlight the value of local ecotypes as reservoirs of unique diversity and as useful case studies for developing pangenome-enabled fingerprinting approaches applicable to varietal protection, conservation, and breeding.

Overall, this study provides a first pangenomic resource for globe artichoke and demonstrates how local ecotypes can serve as informative entry points for integrating comparative genomics, population structure analysis, and molecular fingerprinting. The results support the genomic distinctiveness of ‘Asti Sorì’ within the analysed material, reveal a pangenome organization dominated by conserved gene content but enriched for functionally relevant accessory components, and identify a compact set of candidate SNPs for varietal discrimination. Future work should expand the pangenome to include a broader representation of cultivated globe artichoke, cultivated cardoon, and wild cardoon diversity, and should validate reduced SNP panels across independent collections and technical platforms. Such developments will be essential to transform the resources generated here into robust tools for varietal protection, traceability, conservation, and breeding in *C. cardunculus*.

## 5 CONCLUSIONS

This study developed a first pangenomic framework for globe artichoke, using the Italian local ecotype ‘Asti Sorì’ as an entry point. The pangenome captured a conserved core gene repertoire alongside an accessory component reflecting accession-specific content, with accumulation curves suggesting a tendency toward saturation within the sampled cultivated germplasm. SNP and PAV analyses provided complementary and broadly consistent views of genomic diversity, consistently resolving ‘Asti Sorì’ as a genomically distinct accession within the analyzed collection. SSR loci and accession-specific private variants, while informative in a limited-panel context, proved insufficient or uncertain as robust diagnostic tools for broader varietal discrimination. GBS-derived SNPs resolved well-supported varietal groups across 45 accessions, and a reduced panel of 50 SNPs recapitulated the main population structure of the full dataset within the analyzed germplasm. This compact marker set provides a practical candidate resource for varietal fingerprinting, DUS testing, and the molecular protection of traditional Italian globe artichoke ecotypes. Further validation across independent germplasm collections will be required to establish this panel as a standardized diagnostic assay.

## Supporting information

Supplementary Figure 1. GO enrichment of accessory genome genes. Top enriched BP, MF, and CC terms; bar length indicates gene count and color indica

Supplementary Figure 2. Genome-wide relationships among the analyzed globe artichoke accessions based on SNP and presence/absence variation (PAV) da

## Abbreviations

AFLP: amplified fragment length polymorphism
BH: Benjamini–Hochberg
BP: biological process
CC: cellular component
CTAB: cetyltrimethylammonium bromide
DUS: distinctness, uniformity, and stability
GBS: genotyping-by-sequencing
GO: Gene Ontology
GWAS: genome-wide association study
KEGG: Kyoto Encyclopedia of Genes and Genomes
MF: molecular function
PCA: principal component analysis
PAV: presence/absence variation
RADseq: restriction site-associated DNA sequencing
SNP: single nucleotide polymorphism
SRA: Sequence Read Archive
SSR: simple sequence repeat
SV: structural variation
WGS: whole-genome sequencing.

## SUPPLEMENTAL MATERIAL

### Supplementary Tables

- **Table S1.** Globe artichoke accessions included in the GBS-based diversity panel used for population structure analysis and reduced SNP fingerprinting. For each accession, the corresponding morpho-phenotypic typology is reported according to the classification adopted in previous globe artichoke germplasm studies. Groups include ‘Catanesi’ (CAT), ‘Violetti di Provenza’ (VPR), ‘Violetti di Toscana’ (VTO), ‘Romaneschi’ (ROM), ‘Macau’ (MAC), ‘Spinosi’ (SPI), and ‘Green et al.’ (GEA).
- **Table S2.** Classification of pangenome gene categories. For each of the 37,198 genes identified in the globe artichoke pangenome, the number of accessions in which the gene was detected as present (n_present) and the corresponding pangenome category (core, softcore, shell, or cloud) are reported, as defined in the Materials and Methods. Genes labeled “0” were not detected as present in any of the sampled accessions under the applied presence/absence calling threshold and are reported separately from the four standard pangenome categories.
- **Table S3.** Accession-specific cloud genes and functional annotation. List of cloud genes private to each of the seven core pangenome accessions (’2C’, ‘A41’, ‘Asti Sorì’, ‘C3’, ‘SP’, ‘VS’, ‘VT’), with associated transcript and gene identifiers, predicted protein product, Gene Ontology (GO) terms, and cross-references to the InterPro, RefSeq, UniProtKB/Swiss-Prot, and eggNOG databases.
- **Table S4.** GO enrichment results for accessory and accession-specific gene sets. Gene Ontology enrichment results (Biological Process, Molecular Function, and Cellular Component categories) for the accessory genome (shell and cloud genes combined), the ‘Asti Sorì’-unique and ‘A41’-unique gene sets, and the shell, cloud, and softcore pangenome macro-categories, each tested against the complete pangenome gene background.
- **Table S5.** Performance of random SNP subsampling panels. Summary statistics (best, mean, standard deviation, median, minimum, and maximum Pearson correlation with the pairwise genetic distance matrix derived from the complete SNP dataset) across 100 random replicates, for each of the three tested panel sizes (50, 75, and 100 SNPs).
- **Table S6.** Reduced SNP panels selected for varietal fingerprinting. Genomic coordinates (chromosome or pangenome scaffold, and position) and reference/alternative alleles (REF/ALT) of the single nucleotide polymorphisms (SNPs) composing the best-performing 50-, 75-, and 100-marker panels identified through random subsampling, as described in the Materials and Methods.

### Supplementary Figures

- **Supplementary Figure 1.** GO enrichment of accessory genome genes. Top enriched BP, MF, and CC terms; bar length indicates gene count and color indicates FDR.
- **Supplementary Figure 2.** Genome-wide relationships among the analyzed globe artichoke accessions based on SNP and presence/absence variation (PAV) data. (A) Principal component analysis (PCA) performed using 34,583 pruned SNPs derived from the globe artichoke pangenome. (B) Maximum-likelihood phylogenetic trees inferred from SNP data (top) and presence/absence variation (PAV) data (bottom). Accessions included in the analyses were: ‘Asti Sorì’ (AS), ‘Romanesco C3’ (C3), ‘Violetto di Toscana’ (VT), ‘Violetto di Sicilia’ (VS), ‘Spinoso di Palermo’ (SP), cultivated cardoon ‘Altilis 41’ (A41), and the globe artichoke reference genotype ‘2C’.

## DATA AVAILABILITY

All the data generated in this study are available as Supplementary Material. Sequencing raw data are publicly available and have been deposited in NCBI (PRJNA1481990).

## AUTHORS CONTRIBUTIONS

**E.P.** contributed to conceptualization, data curation, formal analysis, funding acquisition, investigation, methodology, resources, supervision, validation, visualization, writing of the original draft, and review and editing. **E.V.** contributed to data curation, formal analysis, investigation, software, visualization, writing of the original draft, and review and editing. **L.G.** contributed to data curation, formal analysis, software, writing of the original draft, and review and editing. **A.A.** contributed to conceptualization, investigation, methodology, supervision, visualization, writing of the original draft, and review and editing. **C.C.** contributed to conceptualization, investigation, methodology, supervision, visualization, writing of the original draft, and review and editing. **C.Carli** contributed to funding acquisition, resources, supervision, writing of the original draft, and review and editing. **L.B**. contributed to conceptualization, formal analysis, investigation, methodology, software, supervision, visualization, writing of the original draft, and review and editing. **M.M.** contributed to conceptualization, data curation, formal analysis, investigation, methodology, software, supervision, visualization, writing of the original draft, and review and editing.

