## Supplementary figures and images for "A first pangenomic framework for globe artichoke supports SNP-based varietal fingerprinting"

### Supplementary Figure 1. GO enrichment of accessory genome genes. Top enriched BP, MF, and CC terms; bar length indicates gene count and color indica

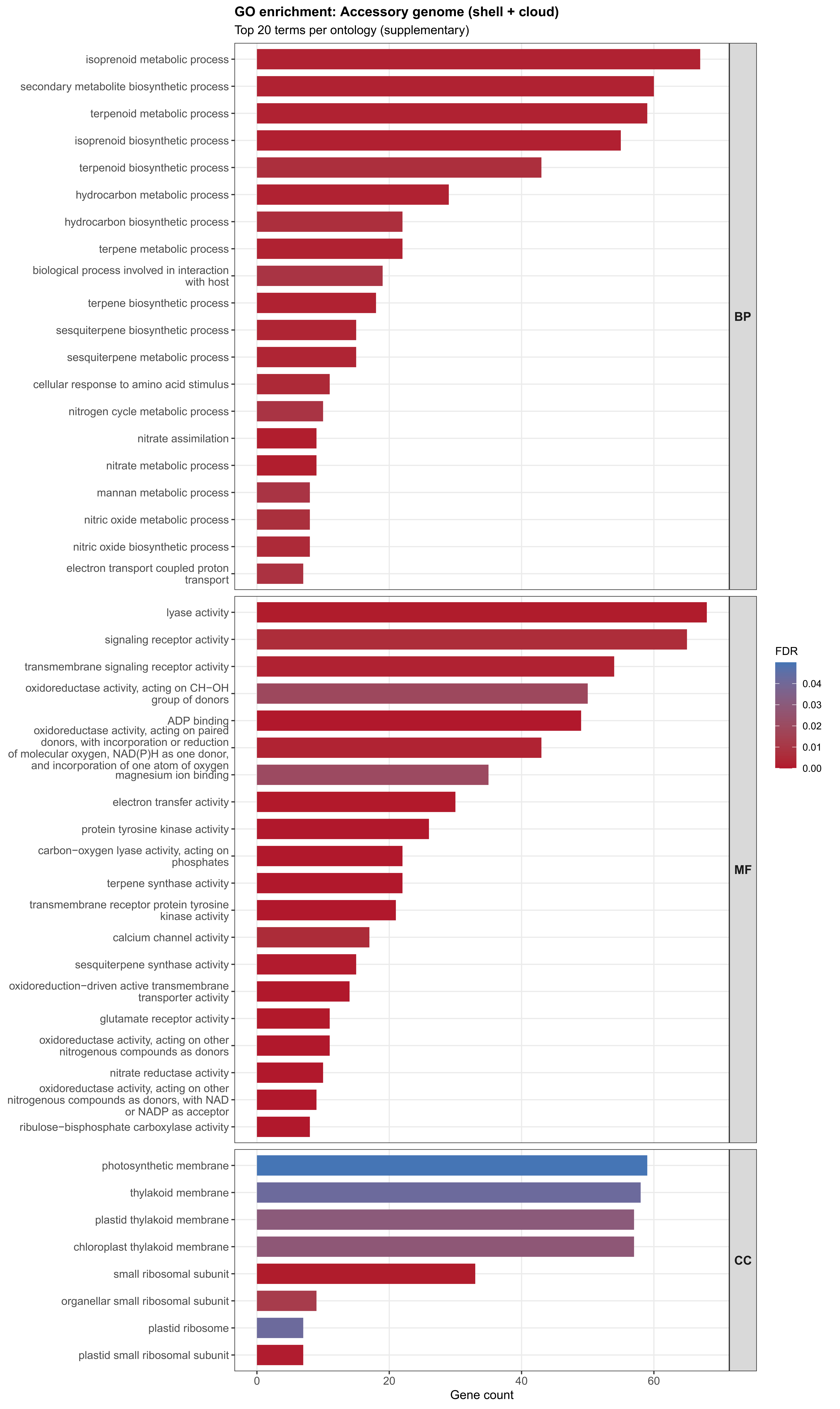

### Supplementary Figure 2. Genome-wide relationships among the analyzed globe artichoke accessions based on SNP and presence/absence variation (PAV) da

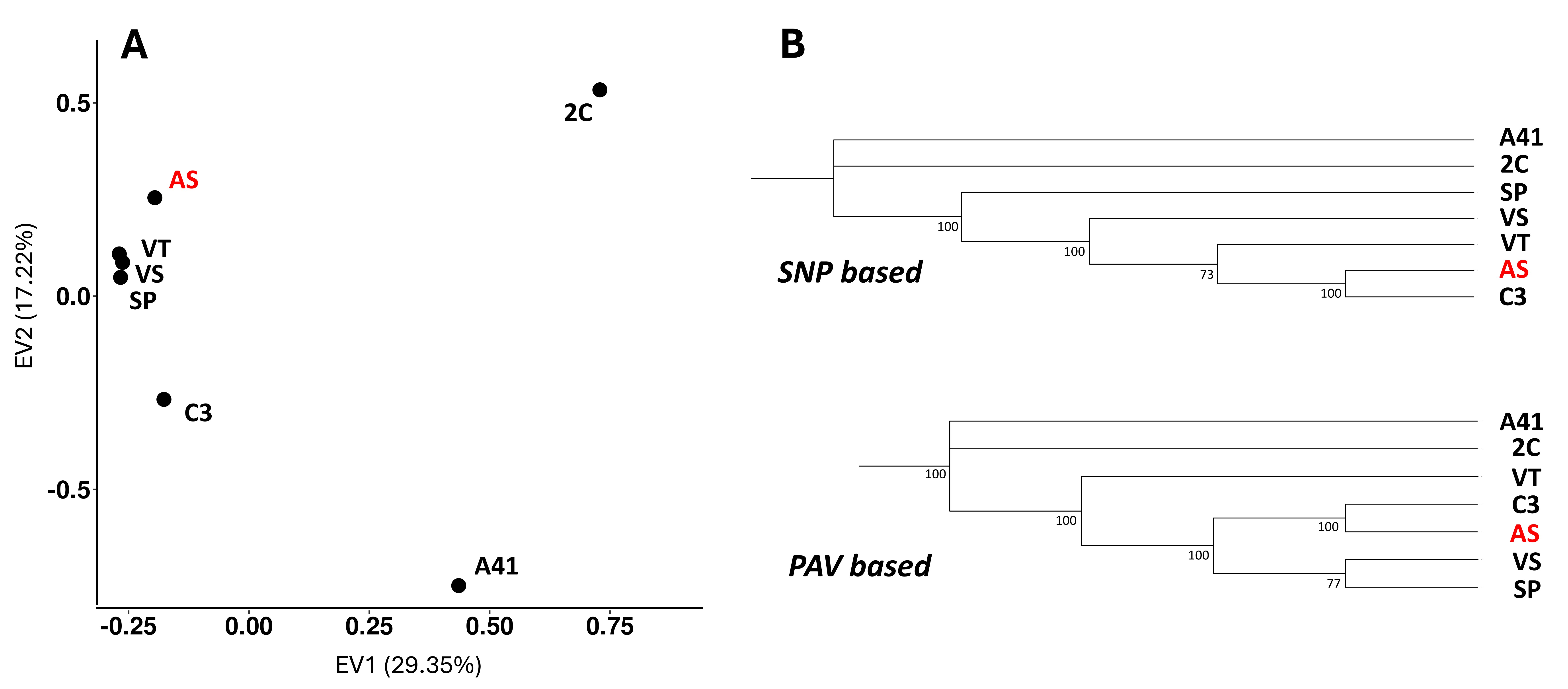
